## Supplemental information for "Phenotypic delay in the evolution of bacterial antibiotic resistance: mechanistic models and their implications"

#### 1 Mutagenesis experiment: mathematical derivations

Here we present derivations of various results discussed in the main text, for the dilution and accumulation mechanisms. The polyploidy mechanism is not discussed here as all results are presented in the main text.

##### 1.1 Dilution of antibiotic-sensitive molecules

In this subsection all cells are assumed monoploid ($c = 1$). Suppose we start from a single genotypically resistant cell containing $n$ sensitive molecules. We follow a single lineage, as in Fig. S1a, and consider the cell in that lineage after $g$ generations. Let the number of sensitive molecules in that cell be $z_n(g)$. A cell is to be considered phenotypically resistant if $z_n(g) = 0$. During cell division, each of the $n$ sensitive molecules may be lost (from the lineage we track) with probability $1/2$. Therefore, the probability that any of the original sensitive molecules present in the initial cell remains in our chosen cell is $2^{-g}$. Hence, $z_n(g)$ is a binomially distributed random variable with $n$ trials and probability parameter $2^{-g}$. The probability of phenotypic resistance in generation $g$ is therefore

$$(1 - 2^{-g})^n \approx e^{-n2^{-g}}. \tag{S1}$$

The exponential approximation in Eq. (S1) holds when $n$ is large and $2^g \propto n$. Resistance thus emerges when $n \approx 2^g$ or for $g \approx \log_2(n)$, in agreement with the simple argument presented in the main text.

We now turn to the probability of resistance occuring in the whole population (for which at least one cell must be resistant), for a population that is initiated with a single mutant cell with $n$ sensitive molecules. Let $p_n(g)$ be the probability that no cell exists with zero sensitive molecules after $g$ generations. Considering the distribution of molecules after the first division we have

$$p_n(g) = \sum_{i=0}^{n} \binom{n}{i} 2^{-n} p_i(g-1) p_{n-i}(g-1). \tag{S2}$$

¶These authors contributed equally to this work.

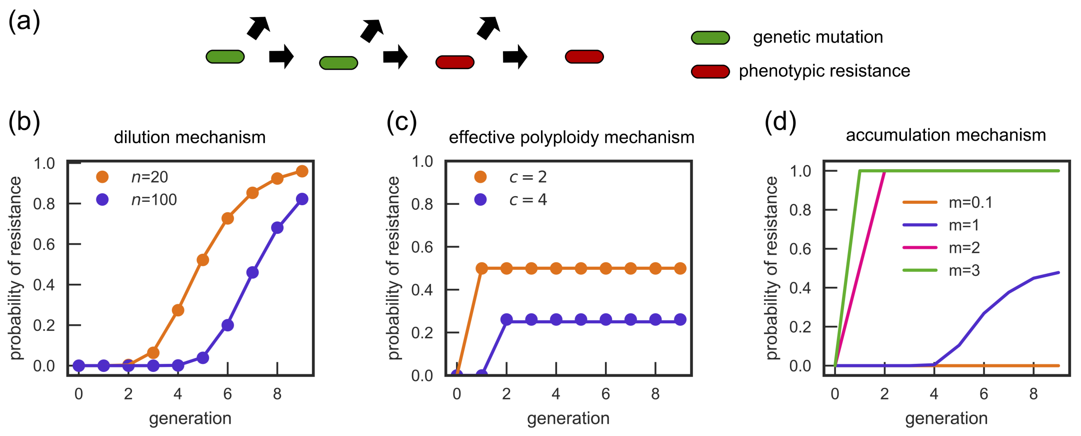


Figure S1: (a) We follow a single bacterium which has just mutated and has the resistant allele in one of its chromosomes. When it divides, we choose one of the two daughter cells at random. After a few generations, this cell can become phenotypically resistant. (b) The probability of the cell being resistant as a function of the number of generations from the genetic mutation for the dilution mechanism (dots: simulation, lines: theory Eq. (S1)). (c) Same as (b) for the effective polyploidy mechanism. (d) Same as (b) for the accumulation mechanism (only simulations).

This recursion allows us to calculate the probability of no resistance after $g$ generations. If we start from $x$ mutant cells rather than just a single mutant, the probability of no resistance after $g$ generations is $(p_n(g))^x$ because mutant lineages are assumed to evolve independently of each other.

To obtain an approximate time for resistance to emerge, we consider the mean number of phenotypically resistant cells. Starting from $x$ cells, after $g$ generations the number of resistant cells can be written as

$$\sum_{i=1}^{x2^g} \eta_i, \tag{S3}$$

where $\eta_i$ is a random variable equal to 1 if the $i$th cell has zero sensitive molecules, and 0 otherwise. The random variables $(\eta_i)_{i=1}^{x2^g}$ are dependent but have identical marginal distributions. Taking expectations and using $\mathbb{E}[\eta_i] = (1 - 2^{-g})^n$, we obtain the expected number of phenotypically resistant cells:

$$\mathbb{E}\left[\sum_{i=1}^{x2^g} \eta_i\right] = x2^g(1 - 2^{-g})^n \approx x2^g e^{-n2^{-g}}. \tag{S4}$$

Let $\tau$ denote the expected time to resistance in the population. From Eq. (S2) the expected time until a resistant cell emerges can be expressed exactly as

$$\mathbb{E}[\tau] = \sum_{k \geq 0} (p_n(k))^x\,, \tag{S5}$$

where $p_n(k)$ can be computed recursively from (S2). In order to obtain an intuitively meaningful result, we reason that phenotypic delay corresponds to the time period during which the expected number of phenotypic mutants is less than 1. Thus a rough approximation for the expected value of $\tau$ is equal to one plus the number of the last generation in which the expected number of resistant cells is less than 1. We use the approximation of (S4) and thus we wish to find $\max\{g : x2^g \exp(-n2^{-g}) < 1\}$. It can be verified by direct substitution that $g = \frac{W(nx) - \log(x)}{\log(2)}$ is the solution, where $W$ is Lambert's function. Approximating $W(z) \approx \log z - \log\log z$ for $z \to \infty$, we obtain that

$$\mathbb{E}[\tau] \approx 1 + \frac{\log n - \log\log xn}{\log(2)} = 1 + \log_2(n/\log(nx)). \tag{S6}$$

Equation (S6) agrees very well with computer simulations (Fig. S2).

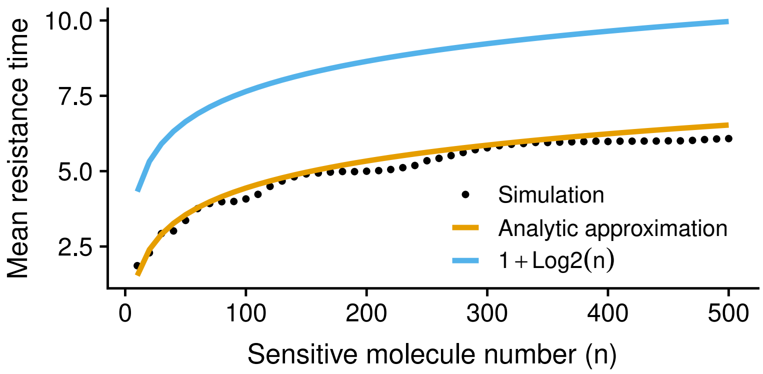


Figure S2: Expected number of generations until a phenotypically resistant cell emerges. We start with $x = 100$ cells that just mutated, and repeat the simulation 500 times for each data point. "Analytic approximation" refers to Eq. (S6).

#### 1.2 Accumulation of resistance-enhancing proteins

We consider the scenario that prior to cell division each genotypically resistant cells create $M_p$ resistance enhancing molecules. These are binomially distributed between the daughter cells at division. A cell is assumed to be phenotypically resistant once it has acquired $M_r$ sensitive molecules.

When tracking a single cell (or random lineage), let $r_g$ be the number of resistant molecules the cell possesses at generation $g$. From the model specification we have the stochastic recursion

$$r_g \stackrel{d}{=} \mathrm{Bin}(M_p, 0.5) + \mathrm{Bin}(r_{g-1}, 0.5), \tag{S7}$$

where $\stackrel{d}{=}$ denotes equality in distribution and we have abused notation using $\mathrm{Bin}(n, p)$ to denote independent binomial random variables with $n$ trials with success probability $p$. Taking expectations over (S7), and solving the resulting recursion, leads to $\mathbb{E}[r_g] = M_p(1 - 2^{-g})$. In fact the full distribution of $r_g$ can be obtained by observing that it is distributionally equal to $\sum_{i=0}^{g-1} \mathrm{Bin}(M_p, 2^{-g+i})$, corresponding to the molecular contributions to our chosen cell from each of the previous generations. This sum has a Poisson-binomial distribution, however the cumulative distribution function (of interest as we care about $\Pr(r_g \geq M_r)$) is relatively uninformative and numerically unstable. Therefore we simply note that the number of molecules after many generations is $M_p$ on average, with unbiased fluctuations around this value (to see that the fluctuations are unbiased set $r_{g-1} = M_p$ in the stochastic recursion). The variance $\mathrm{Var}(r_g) = M_p(2/3 - 2^{-g} + 4^{-g}/3)$ tends to $2M_p/3$ in the limit $g \to \infty$, hence fluctuations become less important for large $M_p$ (the coefficient of variation tends to zero).

As a first approximation, since our condition for resistance is that $\mathbb{E}[r_g] \geq M_r$, then resistance will occur at $\tau \approx -\log_2(1 - 1/m)$ as long as $m = M_p/M_r > 1$. Note that if $m < 1$, $r_g$ may stray above $M_r$ due to fluctuations, but this cannot produce sustained resistance as such high $r_g$ values will be transient. We therefore do not consider the case of $m < 1$ in detail. We also omit the special case $m = 1$ for lack of biological realism.

Turning to the population as whole, but still starting with a single cell, we firstly note that, as with Eq. (S2), the following recursion holds for the probability of no resistance after $g$ generations starting with $n$ resistance molecules:

$$p_n(g) = \sum_{k=0}^{M_p+n} \binom{n + M_p}{k} 2^{-n-M_p} p_k(g-1) p_{n+M_p-k}(g-1). \tag{S8}$$

Initial conditions for the recursion are $p_n(0) = 1$ for $n < M_r$, and 0 otherwise. Again, for $x > 1$, the probability of no resistance is $(p_n(g))^x$.

In summary, for $m < 1$ resistance will not be stably achieved (steady state less than $M_r$), while if we seek a delay of at least a generation ($\tau \geq 1$) we require $m \leq 2$. A noticeable phenotypic delay occurs thus only in the narrow parameter range $1 \leq m \leq 2$. As mentioned in the main text, resistance will occur faster in the whole population than down any lineage, further narrowing the parameter regime in that setting.

##### 1.3 Combining the dilution and polyploidy mechanism: phenotypic penetrance

We now consider bacteria with ploidy $c$. Starting from a cell possessing a mutated allele on a single chromosome, $g_c = \log_2(c)$ divisions are required to generate a cell with all $c$ copies having the resistant allele (henceforth termed chromosomal resistant). To include the dilution mechanism, suppose $n$ sensitive molecules exist in the initial cell and that each wild-type chromosome produces $n/c$ sensitive molecules between cell divisions.

A descendant of the initial mutated cell that emerges with a full suite of resistant chromosomes will eventually have phenotypically resistant cells amongst their progeny (after dilution of any sensitive molecules). This cell will initially have $n_c$ sensitive molecules. Binomial partitioning of the $n$ original sensitive molecules and those created between the appearance of the first mutant choromosome and the chromosomally resistant cell then allows us to write

$$n_c \overset{d}{=} \mathrm{Bin}(n, 2^{-g_c}) + \sum_{i=0}^{g_c-1} \mathrm{Bin}\left((1-2^i/c)n, 2^{-(g_c-i)}\right) \tag{S9}$$

where Bin(...) denotes, as before, independent binomial random numbers. Note that here we restrict ourselves to $n$ such that $(1-2^i/c)n$ is an integer, for each $0 \le i \le g_c - 1$. This is satisfied if each sensitive chromosome produces an integer number of sensitive molecules ($n/c$).

Any phenotypically resistant cells will be descendants of the initial chromosomally resistant cell. The question of whether there is any such resistant cell by generation $g$ is therefore equivalent to asking whether there is any resistant cell by generation $g - g_c$ but initiating the process with the initial chromosomally resistant bacteria with $n_c$ sensitive molecules. To find the expected number of phenotypically resistant cells, we average over (S4) with respect to the distribution of $n_c$, that is we compute

$$\mathbb{E}[2^{g-g_c}(1-2^{-(g-g_c)})^{n_c}] = 2^{g-g_c}\mathbb{E}[(1-2^{-(g-g_c)})^{\mathrm{Bin}(n,2^{-g_c})+\sum_{i=0}^{g_c-1}\mathrm{Bin}((1-2^i/c)n,2^{-(g_c-i)})}] \tag{S10}$$

$$= 2^{g-g_c}(1-2^{-g})^n \prod_{i=0}^{g_c-1}(1-2^{-(g-i)})^{n(1-2^i/c)}. \tag{S11}$$

Here, the generating function for the binomial distribution has been used. If, as before, we define genotypically resistant cells as having at least one resistant chromosome, then for any $g \le g_c$ only a single genotypically resistant bacterium will exist because we have assumed co-inheritance of recently linked chromosomes. After generation $g_c$ all genotypically resistant cells are the descendants of the initial chromosomally resistant cell, and their number is $2^{g-g_c}$. Hence, of the genotypically resistant cells, the expected fraction of phenotypically resistant cells is given by

$$\begin{cases} 0 & 0 \le g < g_c, \\ (1-2^{-g})^n \prod_{i=0}^{g_c-1}(1-2^{-(g-i)})^{n(1-2^i/c)} & g_c \le g. \end{cases} \tag{S12}$$

which gives Eq. (1) in the main text. Following [1] we call this quantity the phenotypic penetrance. The limiting case of $n = 0$ gives a Heaviside step function $\theta(g - g_c)$ as expected. Note that while this equation was derived by considering the scenario where we start with a single mutated cell, it holds also when there are initially many mutated cells ($x > 1$). This is because phenotypic penetrance is the ratio of phenotypically resistant cells to genotypically resistant cells.

### 2 Single-cell simulations

One of the key arguments used by Sun et al. [1] to affirm that phenotypic delay is caused mostly by the effective polyploidy mechanism was the asymmetric inheritance of resistance, i.e., the observation that some lineages become resistant while others do not. This effect can only be observed by studying the behaviour of individual cells. Hence, to understand the effect of the different phenotypic delay mechanisms, we simulate a single cell immediately after a mutation (figure S1a). When the cell divides, we follow one randomly selected daughter and continue tracking that cell until phenotypic resistance emerges or the cell loses all resistant alleles. We repeat the simulation 10000 times and calculate the probability of the cell becoming phenotypically resistant as a function of the number of generations from the mutation.

In the case of the dilution mechanism, the phenotypic delay is controlled by the number of molecules $n$ that need to be diluted out (figure S1b). More molecules lead to a longer delay because more generations must pass before random segregation produces a cell without any sensitive molecules. In contrast to the effective polyploidy model, the probability of phenotypic resistance always approaches 1 for long times, and the approach to this end point is also more gradual. This is because, unlike in the polyploidy model, molecules segregate independently of each other, and the probability of producing a cell with only resistant molecules is non zero (albeit very small) already after the first division. Since genetic resistance cannot be lost in this model (there is no chromosomal segregation), all cells eventually become resistant. Analytically, we showed (equation S1) that the probability of resistance is equal to :

$$P_{\text{res}}(g) = (1 - 2^{-g})^n, \tag{S13}$$

where g is the number of generations from the mutation.

In the case of the effective polyploidy mechanism, the probability of resistance is exactly zero until generation $1 + \log_2 c$, after which it assumes a constant value $1/c$ (figure S1c), where $c$ is the effective polyploidy. This is because if only one of the $c = 1, 2, 4, \ldots$ chromosomes has the resistant allele, exactly one cell out of $c$ progeny of the initial cells ends up with resistant chromosomes, and the rest $c - 1$ cells have only sensitive chromosomes. Therefore,

$$P_{\text{res}}(g) = \begin{cases} 0 & \text{if } g < 1 + \log_2 c \\ 1/c & \text{if } g \geq 1 + \log_2 c. \end{cases} \tag{S14}$$

In figure S1c, we observe close agreement between the theory and our stochastic simulations.

The third mechanism that could lead to a phenotypic delay is the accumulation of resistance-enhancing molecules (figure S1d). As explained in the main text and in Section 1.2, a substantial phenotypic delay is only observed within a narrow parameter range for this mechanism: $1 \lesssim m \lesssim 2$.

### 3 Biasing the partitioning of molecules at divisions shortens phenotypic delay

In the main text, for the dilution mechanism we assumed that target molecules are divided between the two daughters without any bias. While this is a reasonable assumption for cytoplasmic proteins, there is evidence that membrane-associated proteins are segregated in a biased way: a fraction $p > 1/2$ ends in a daughter cell with the older pole [2, 3, 4]. In particular, a bias $p = 0.62$ has been shown for efflux pumps [5], whose expression is associated with elevated resistance to antibiotics. Another example is OmpA (outer membrane protein A) which also tends to accumulate in cells with older poles [3]. OmpA is implicated in phage invasion; mutations that decrease or alter the protein create resistance to bacteriophages [6, 7].

To investigate the effect of biased protein segregation on the length of phenotypic delay, we repeated our single-cell and population level simulations using the dilution mechanism ($n = 1000$) with a biased binomial distribution ($p = 0.62$ instead of $p = 0.5$ as used in the main text). A very small effect of the bias is present at the single-cell level (figure S3a), but a much larger effect (delay shortened by one generation) is observed at the population level (figure S3b). This can be explained as follows. At the population level, it is enough for a single cell to become resistant. Biasing causes cells with the younger pole to be depleted of the sensitive molecules faster and, hence, to become resistant earlier. On the other hand, at the single-cell level we follow a randomly selected daughter cell and not necessary the one with the younger pole. This largely nullifies the effect because even though some cells in the lineage get rid of the sensitive molecules faster than in the case of no bias, others take longer.

### 4 Phenotypic delay when the number of target molecules is independent of the growth rate

In the main text, for the combined dilution and effective polyploidy mechanisms we assumed that the number of sensitive target molecules depends on the growth rate. In particular, we assumed that the gyrase enzyme constitutes a roughly constant fraction of the proteome, independently of

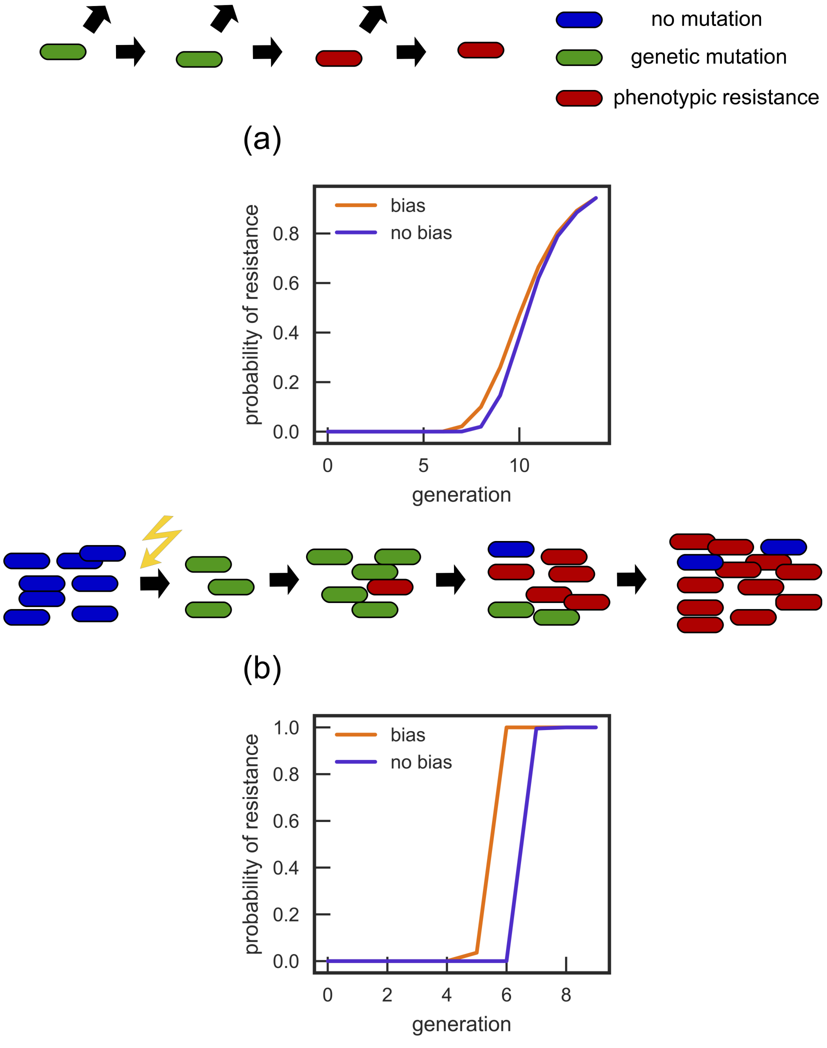


Figure S3: Biasing the segregation of sensitive molecules at division leads to a decrease in the phenotypic delay both at the (a) single-cell and (b) population level. Blue curve represents an unbiased case ($p = 0.5$), orange curves is the biased case ($p = 0.62$). In all cases, $n = 1000$.

the growth rate [8]. Assuming that the total protein mass is proportional to cell volume, and that volume $\propto 2^{\lambda/\lambda_0}$, where $\lambda$ is the growth rate and $\lambda_0 = 1\text{h}^{-1}$ [9, 10], we can then predict the change in the number of gyrase molecules per cell associated with a change in growth rate. For example, if we assume that $n = 20$ for $t_d = 60\,\text{min}$, then for $t_d = 30\,\text{min}$ we obtain $n = 40$ (figure S4a). In contrast, in this subsection we investigate what happens if the number of target molecules is independent of the doubling time. To be specific, let us take $n = 20$ for both $t_d = 60\,\text{min}$ and $t_d = 30\,\text{min}$ (figure S4b). We observe that when the number of target molecules $n$ does not depend on $t_d$, the increase in phenotypic delay decreases from 2 to 1 generation, similar to the results obtained for the effective polyploidy model without the dilution mechanism. Hence, the increase in phenotypic delay in the combined model caused by a decrease of the doubling time $t_d$ is caused largely by the increase in the number of target molecules.

In the main text, we also show that the probability of surviving an antibiotic treatment in a simulated infection for the combined model is a function $t_d$ (figure S4c). In figure S4d we compare this to the case where the number of sensitive target molecules $n$ does not depend on $t_d$. An almost complete lack of dependence on $t_d$ is observed in this case, showing that the dependence on the doubling time seen in figure S4c is again mostly due to the change in $n$ with doubling time.

### 5 Phenotypic delay for the dilution mechanism with non-zero molecular threshold for resistance

In the main text, for the dilution mechanism, we assumed that a cell becomes resistant only when it loses all $n$ sensitive molecules. However, in reality a cell may be resistant even if a few sensitive molecules are left. Here we repeat the analysis for the dilution mechanism for which the threshold in the number of sensitive molecules for which a cell is phenotypically resistant is $n_\text{r} > 0$. The results are summarized in figure S5. As expected, the phenotypic delay decreases as $n_r$ increases, because the number of molecules required to be diluted out for resistance to emerge decreases. Interestingly, we also observe that increasing $n_r$ leads to a less gradual appearance of resistance at the single-cell level.

### 6 Maximum likelihood estimate of the mutation rate

In figure S6 we show the maximum likelihood estimates for the mutation probability, obtained using the package flan [11], from 1000 simulations of the fluctuation test from Ref. [12]. Simulations were performed using the same method as discussed in the main text, Section 4.6. Each of the 1000 simulated experiments contained 40 independent replicates. Replicates were initiated with $N_0 = 100$ cells and the final population size was $N_f = 2^{20} N_0$ so that the population of bacteria grew for 20 generations. A relatively narrow range of estimated mutation probabilities was obtained, as shown in figure S6. This strongly suggests that the observed discrepancy between the mutation rates obtained from sequencing and the fluctuation test is not due to sampling variation but is likely to have another (biological) cause. We suggest that this may be phenotypic delay.

### 7 Sensitivity analysis of the ABC method of model comparison

In the main text we report that the probability that the data of Boe et al. [13] is generated by the phenotypic delay model with dilution, compared to the model with no delay, is 0.97.

To check how robust this result is, we performed two sensitivity analyses, altering the size of our simulation bank $N_\text{sim}$, and the number of 'closest' simulations we compare to, $N_\text{thresh}$ ($N_\text{thresh} = 100$ in the main text). Firstly, we query whether enough simulations have been performed. To do so we sub-sample our bank of simulations. Secondly, we assess how our probability estimate changes as a function of $N_\text{thresh}$. Note that as $N_\text{thresh} \to N_\text{sim}$ the probability estimate tends to 0.5 as an equal proportion of simulations comes from either model. The results are presented in Fig. S7, which supports our conclusion that the model with delay via the dilution mechanism is indeed a better generative model for the data than the model without delay.

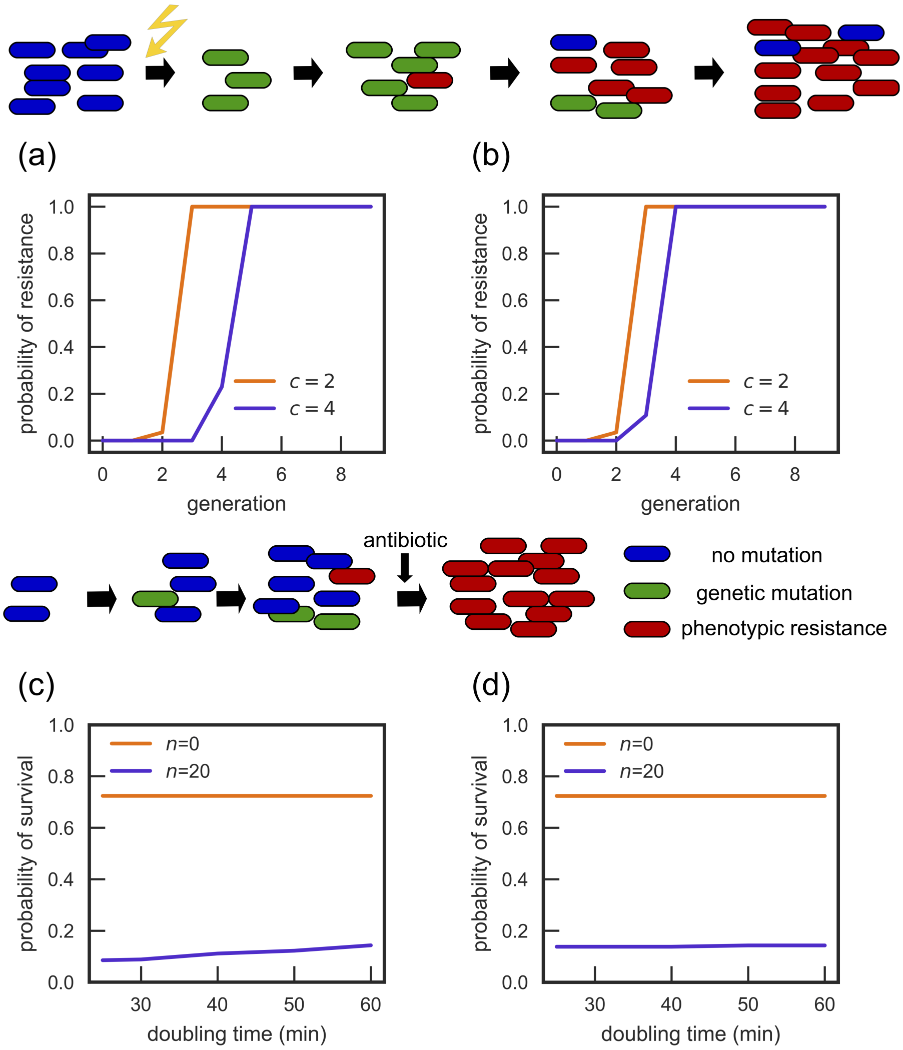


Figure S4: Effect of dependence of the number of target molecules on the doubling time $t_d$ for the combined model. (a) Probability of resistance as a function of time (generations) for different doubling times (determined by ploidy $c$) when the number of target molecules $n$ depends on $t_d$. (b) Same as (a) but for the model in which $n$ does not depend on $t_d$. (c) Probability of survival for a simulated infection (see section 2.3 and figure 3 in the main text) for a combined model when the number of target molecules depends on the growth rate. (d) Same as (c) but for the model in which $n$ does not depend on $t_d$.

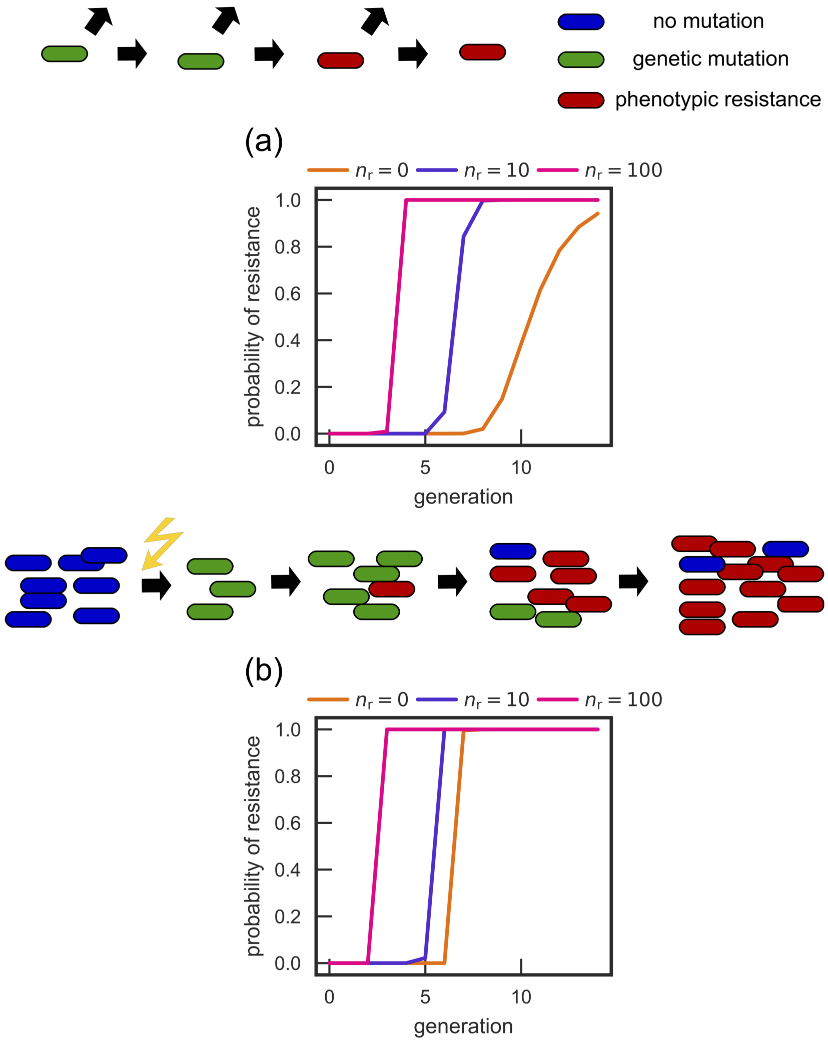


Figure S5: A partial dilution mechanism decreases the phenotypic delay. (a) Single-cell and (b) population level simulated experiments as a function of $n_r$, the number of sensitive molecules allowed for resistance to emerge. In all cases, the total number of molecules $n = 1000$.

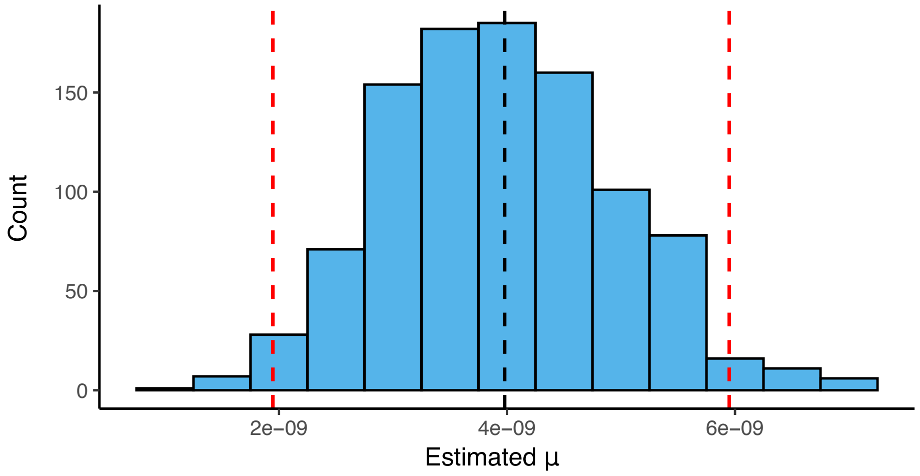


Figure S6: Maximum likelihood estimates of $\mu$ from 1000 simulations mimicking the experiment of Ref. [12] with known $\mu = 3.98 \times 10^{-9}$ (black vertical line) for the no-delay model. The mutation probability can be underestimated by a factor of 2 (95% of simulations yielded estimates between red vertical lines), whereas Ref. [12] reports a factor of 9.5 difference between $\mu$ obtained from DNA sequencing and fluctuation tests. The Lee et al. result [12] cannot be thus explained by the no-delay model.

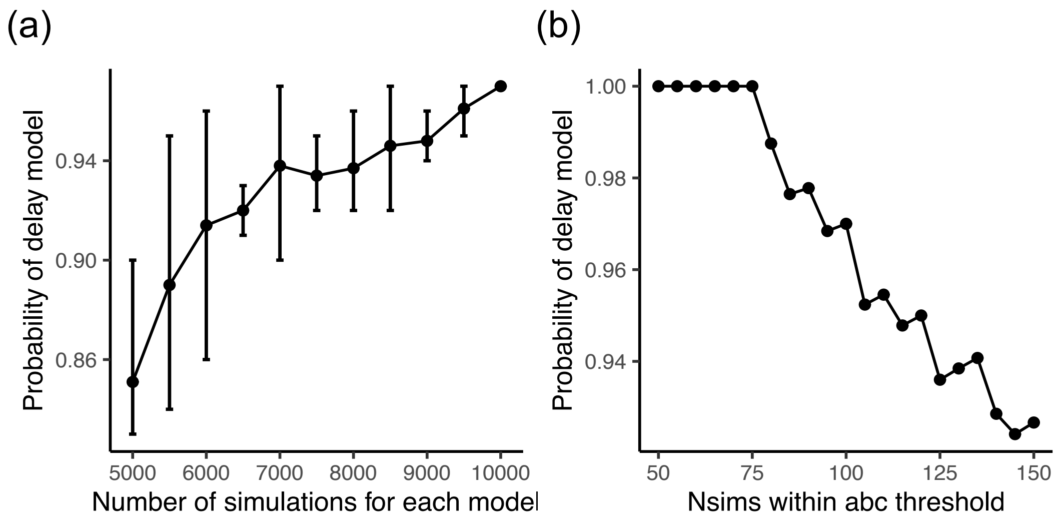


Figure S7: Sensitivity analysis for model selection. (a) The probability of the Boe et al. data [13] coming from the delay model as a function of the number of simulation runs. The runs were randomly sampled from the original bank of simulations and the probability of the delay model was estimated. The process was repeated 10 times. Error bars are the maximum and minimum probability estimated, with the centred dot as the mean. (b) The probability estimate for the probability of the data [13] coming from the delay model as a function of $N_{\text{thresh}}$.

336. doi:10.1038/nature14461.
